# Preparatory activity links frontal eye field activity with small amplitude motor unit recruitment of neck muscles during gaze planning

**DOI:** 10.1101/2021.02.01.429125

**Authors:** Satya P. Rungta, Debaleena Basu, Naveen Sendhilnathan, Aditya Murthy

**Author notes:** **Corresponding author:** Prof. Aditya Murthy, Centre for Neuroscience, Indian Institute of Science, Bengaluru – 576104, Karnataka, India, **Email:**.

## Abstract

A hallmark of intelligent behavior is that we can separate intention from action. To understand the mechanism that gates the flow of information between motor planning and execution, we compared the activity of frontal eye field neurons with motor unit activity from neck muscles in the presence of an intervening delay period in which spatial information regarding the target was available to plan a response. Whereas we could infer spatially-specific delayed period activity from the activity of frontal eye field neurons, neck motor unit activity during the delay period could not be used to infer the direction of an upcoming movement, Nonetheless, motor unit activity was correlated with the time it took to initiate saccades. Interestingly, we observed a heterogeneity of responses amongst motor units, such that only units with smaller amplitudes showed a clear modulation during the delay period. These small amplitude motor units also had higher spontaneous activity compared to the units which showed modulation only during the movement epoch. Taken together, our results suggest that the temporal information is visible in the periphery amongst smaller motor units during eye movement planning and explains how the delay period primes muscle activity leading to faster reaction times.

**Significance statement:** This study shows that the temporal aspects of a motor plan in the oculomotor circuitry can be accessed by peripheral neck muscles hundreds of milliseconds prior to the instruction to initiate a saccadic eye movement. The coupling between central and peripheral processes during the delay time is mediated by the recruitment pattern of motor units with smaller amplitude in the periphery. Besides giving insight into how information processed in cortical areas is read out by the muscles, these findings could be useful to decode intentional signals from the periphery to control brain machine interface devices.

## Introduction

A hallmark of intelligent behavior is that we can couple or decouple our intentions from actions. To enable such flexibility, the nervous system has evolved multiple possible mechanisms to gate the flow of information between the central nervous system and the peripheral musculature. For example, only certain patterns of activity in the motor cortex, that are associated with movements, are allowed to flow into the periphery while other patterns of activity can process information locally and cancel out each other thereby preventing the flow of information into the periphery(1). The basal ganglia are thought to be one of the structures that gate the flow of information between the cortex and the spinal cord via strong inhibitory control (2–7) Additionally, local processing of information involving inhibitory interneurons within the brainstem and spinal cord circuitry (8–10) are also thought to play an important role in decoupling signals in the spine that carry information about motor plans (8, 9, 11) and motor decisions (12) from the peripheral musculature.

Despite the presence of gating mechanisms, certain behavioral paradigms, which facilitate the rapid planning and execution of visually-guided saccadic eye movements (13), have shown that stimulus-related activity can be elicited from neck muscles of head-fixed monkeys despite the absence of overt eye movements (14–16). Additionally, a visual target is known to initiate reflexive recruitment of proximal limb muscles in cats, monkeys and humans (17–20), and stimulus-linked activity can also be recorded from shoulder muscles during such rapid reflexive movements but not during delayed movements (21), suggesting that muscle activity is gated for deliberative movement that require preparation (22–26). However, in contrast to the dominant view, the presence of delayed period motor unit activity in axial and proximal skeletal muscles during a delayed movement task in humans and monkeys has been shown (27, 28).

In light of these observations, in this study, we tested whether the delay period activity can be elicited from neck muscles, and whether this activity parallels the activity in frontal eye field neurons (FEF) that convey gaze-related signals to the neck muscles. We tested whether such delay period activity can be modelled in a manner analogous to accumulator models, along with their implication on behavioral reaction times. For this, we simultaneously recorded motor units from the neck muscles and neurons from FEF, during memory-guided saccade behavior in which the delay period separated ‘where’ to look from ‘when’ to initiate a movement. This separability allows one to characterize and model the nature of the delay period activity in neck muscles and test different aspects of a motor plan that is transmitted to the periphery in the absence of overt movements.

## Results

Two monkeys performed a memory-guided saccade task (**Fig 1A**) in which the saccade target was briefly flashed in any one of the eight possible locations, followed by a delay period of ~1000 ms, after which the monkeys had to make a saccade to the remembered target location to earn a liquid reward (see methods for details). The memory-guided saccade task requires successful encoding of the target stimulus, maintaining the location in memory during the delay period, and then making a saccade to the remembered location, thus allowing for separation of visual, delay, and movement related processes. We recorded 210 FEF neurons from two monkeys and 152 motor units from dorsal neck muscles (splenius capitis) simultaneously as the monkeys were performing the task (**Fig 1B**).

**Figure 1:**
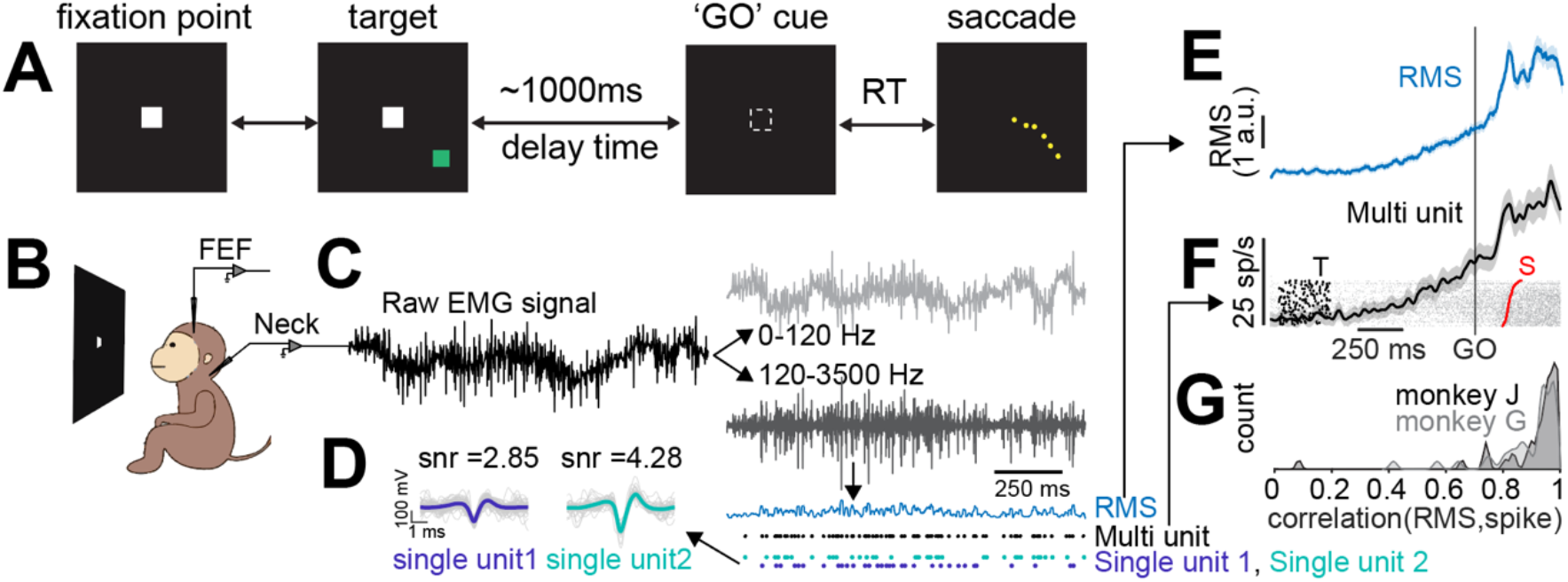
Task and method. **A)** Memory guided saccade task paradigm **B)** Schematic of the recording setup **C)** Raw data that was collected from the muscle EMG was low-pass filtered and high-pass filtered. Windowing and thresholding were used to calculate root mean square (RMS) and detect putative multi units respectively. The multi-unit activity was then separated into multiple single units based on waveform amplitude. **D)** Two single unit waveforms with their signal to noise ratio (snr) isolated from the same multi-unit. **E)** Representative muscle activity from a session, aligned to time of go cue, calculated by using the conventional approach of average root mean square (RMS) of the processed signal **F)** Same muscle activity from above but calculated using a raster based method. Gray markers are individual spikes, black markers are time of target onset (T) and red markers are time of saccade onset (S). **G)** Density histogram showing the Pearson’s correlation coefficient for data analyzed using the two approaches all sessions (Correlation between **Figs 1E** and **1F** for all sessions) for monkey J (in black) and monkey G (in gray).

We analyzed the neural signal based on traditional methods: by convolving the spike timings with a temporal Gaussian filter and obtaining a spike density function (see methods). However, we analyzed the EMG activity through two distinct methods: (a) the traditional root mean square approach and (b) the same signal processing techniques we used for the neural data. In the former approach, the envelope or variability in the EMG signal was captured using a running window over the signal squared, whereas in the latter approach, EMG signals were first decomposed into multi-unit and single unit activities (see methods, **Fig 1C**) and after isolating single unit spike waveforms (**Fig 1D**), spike timings were convolved using a temporal Gaussian filter and obtaining a spike density function (**Fig 1F;** see methods). Both the approaches gave comparable results (**Fig 1G;** Monkey J: Pearson’s r = 0.89 ± 0.02, p<0.0001; Monkey G: Pearson’s r = 0.88 ± 0.03, p<0.0001; Both: 0.88 ± 0.03; p<0.0001). However, since the latter method provided information about the spike waveform in addition to the frequency of spike occurrence, and was the same method we used for analyzing the neural data, we only used the latter analyses henceforth, unless otherwise specified.

We typically isolated 1 to 2 single motor units (maximum of 3) from each multi-unit intramuscular recordings per session. For example, from the average EMG recording shown in **Fig 2A top panel**, we could isolate two different motor units shown in **Fig 2A bottom panel**. Both the motor units showed two different kinds of responses. We called the unit that showed ramping up of activity, prior to the go cue as ‘rampers’ and the unit that showed no change in activity during the delay period as ‘non-rampers’.

**Figure 2:**
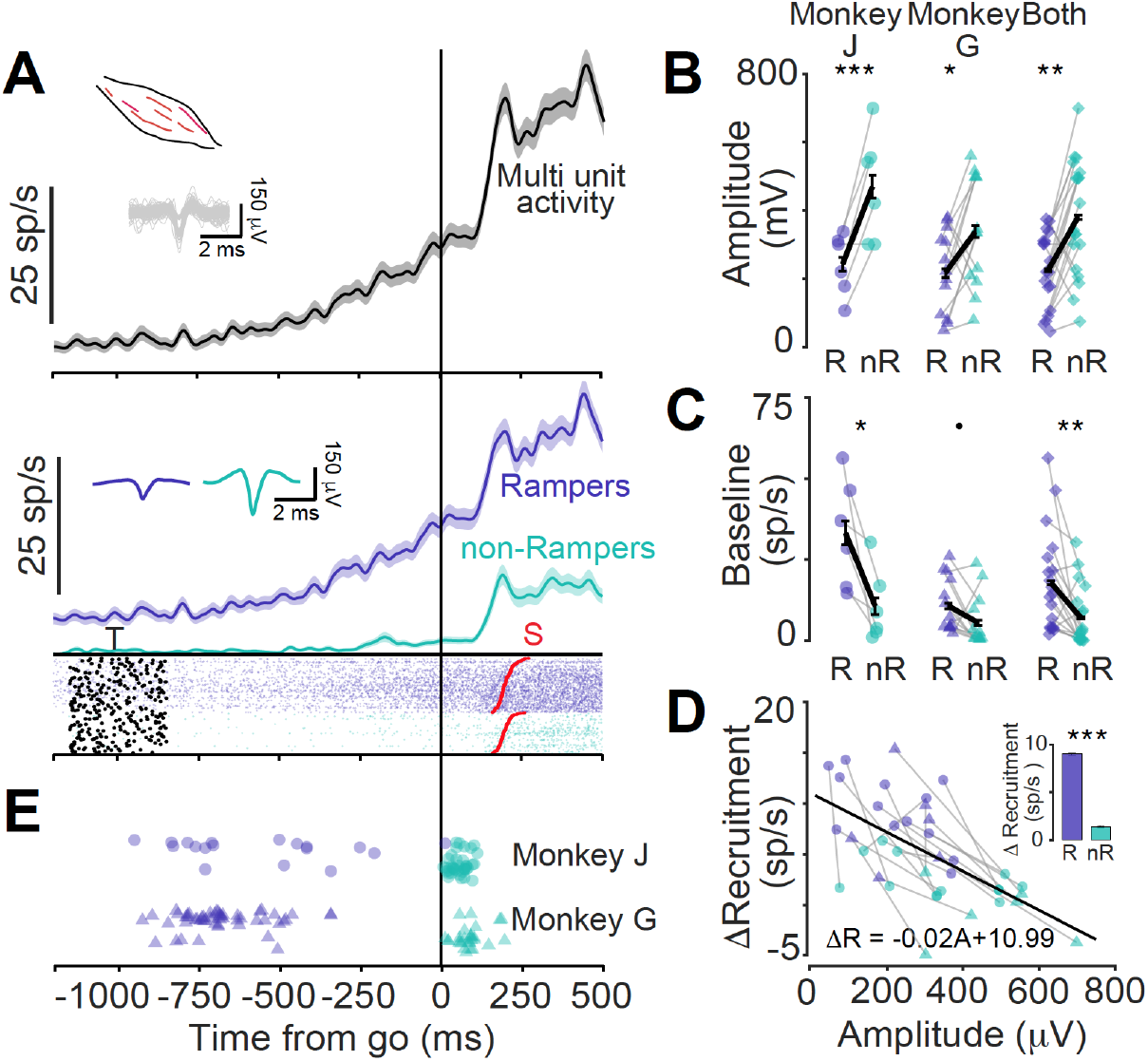
Ramper and non-ramper motor units. **A)** Top: Multi unit activity from a representative session. Inset shows the spike waveforms. Bottom: activity for two motor units that were isolated from the session (Purple: rampers, Cyan: non-rampers). Raster same format as **Fig 1F**. **B)** Average amplitude of spike waveforms for rampers (purple) and non-rampers (cyan) for monkey J (left), monkey G (center) and combined (right). * means p<0.05; ** means p<0.01; *** means p<0.001. **C)** Average basal firing rate for rampers (purple) and non-rampers (cyan) for monkey J (left), monkey G (center) and combined (right). * means p<0.05; ** means p<0.01 **D)** Recruitment pattern for rampers (purple) and non-rampers (cyan). Each gray line represents the data collected during a single session for monkey J (circles) and monkey G (triangles). The black line represents the best fit linear regression model for the data set. The bar plots in the inset represent the average recruitment for the rampers and non-rampers at the time of go cue. *** means p<0.001 **E)** Time of onset for rampers (purple) and non-rampers (cyan) for monkey J (circles) and monkey G (triangles).

These two types of motor units had several different functional characteristics: First, rampers had significantly smaller spike waveform amplitudes (Monkey J: 241.92 ± 36.7 μV, Monkey G: 216.514 ± 33.30 μV, both: 224.538 ± 25.83 μV) than non-rampers that were recorded simultaneously during the same session (Monkey J: 469.53 ± 64.33 μV, Monkey G: 337.03 ± 44.83 μV, Both: 378.87 ± 38.60 μV, Pairwise t-test: Monkey J: t(5)=-4.34; p = 0.007; Monkey G: t(12)=2.39; p = 0.034; For both: t(18)=-3.93; p = 0.01; **Fig 2B**). Second, rampers had a higher baseline firing rate (Monkey J: 33.23 ± 6.77 sp/s, Monkey G: 10.67 ± 2.33 sp/s, both: 17.79 ± 3.55 sp/s) compared to non-rampers (Monkey J: 10.51 ± 4.61 sp/s, Monkey G: 5.42 ± 2.24 sp/s, both: 7.03 ± 2.12 sp/s; Pairwise t-test: Monkey J: t(5)=2.98; p=0.031; Monkey G: t(12)=1.87; p=0.086; For both: t(18)=3.06; p=0.007; **Fig 2C**). Third, changes in the recruitment pattern was higher in rampers (Monkey J: 7.93 ± 1.82 sp/s, Monkey G: 9.48 ± 0.97 sp/s, both: 8.99 ± 0.86 sp/s) than non-rampers (Monkey J: −.82 ± 1.23 sp/s, Monkey G: 2.45 ± 0.54 sp/s, both: 1.41 ± 0.62 sp/s; Pairwise t-test: Monkey J: t(5)=-5.78; p=0.002; Monkey G: t(12)=-5.18; p<0.001; For both: t(18)=-7.30; p<0.001; combined result shown as inset in **Fig 2D**). Furthermore, the change in activity during delay time was negatively correlated (Linear regression: β=-0.02, p<0.001; Pearson’s r = −0.61, p<0.001) to the amplitude size of the wave form isolated and captured during the same session. **Fig 2D**). Also, we noticed that across multiple sessions, the strength of the modulation during the delay time was not correlated to the baseline firing activity for both rampers (Pearson’s r = 0.097, p = 0.693) as well as non-rampers (Pearson’s r = −0.058, p = 0.813). Finally, rampers were recruited much earlier during the delay period (Monkey J: −564.72 ± 61.52 ms, Monkey G: −683.17 ± 18.17 ms; For both: – 652.71 ± 42.35 sp/s), whereas non-rampers were recruited much later after the go cue (Monkey J: +49.89 ± 5.51 ms; Monkey G: +86.41 ± 9.65 ms; For both: 60.47 ± 6.74 sp/s; **Fig 2E**). Also, the rampers (Monkey J: 24.51 ± 6.50 sp/s, Monkey G: 20.88 ± 2.70 sp/s; For both: 21.82 ± 3.99 sp/s), had higher slope compared to non rampers (Monkey J: 0.05 ± .73 sp/s, Monkey G: 1.39 ± 1.77 sp/s; For both: 0.60 ± .75 sp/s).

In summary, motor units that were recruited early during the delay period, the rampers, had smaller amplitudes, higher spontaneous and peak firing activity compared to units that were recruited later, the non-rampers. We only further examined rampers in this study. A significant number of motor units (46.05 %, 70 out of 152) and neurons (44.28 %, 93 out of 210) showed modulations during the delay epoch. Therefore, we next asked what information did these motor units encode in the delay epoch. To study this, we looked at the spatial and temporal information encoded in the motor units and compared them with that of the FEF neurons as discussed below.

### Spatial information about saccade plan during the delay period is available in the FEF neurons but not in neck muscle motor units

We first asked whether the ramper motor unit activity during the delay period contained any spatial information about the upcoming saccade. We characterized each motor unit’s response field (RF) as the direction with maximum activity, −125 to 25 ms relative to saccade onset (**Fig S2A**). Different motor units had different preferred directions showing a broad range of selectivity in their responses. The average directional tuning across the population for right (Monkey J: 42^0^, Monkey G: 48^0^) and left musculature (Monkey J: 153^0^, Monkey G: 137^0^), indicated a horizontal and upward component in the composite motor unit activity (**Fig S2B**).

To calculate the time when each unit could reasonably predict the direction of an upcoming saccade, we first divided the trials into 2 groups: trials where saccade went into the RF (in-RF) and trials where the saccades went outside the RF (out-RF). We then performed an ROC analysis on the EMG and the neural activity between in-RF and out-RF responses, for visual, delay and movement epochs and calculated the area under the curve statistic (AUC). AUC values range between 0.5 (no separability) to 1 (complete separability). Even though motor units showed modulation during the delay period, the responses of the units for in-RF and out-RF conditions, during the delta epoch, were comparable (**Fig 3A**) unlike neurons (**Fig 3B**). At the population level, the discriminability between in-RF and out-RF responses increased continuously with time, from target onset to movement onset, for neural activity but remained largely unchanged for EMG until the go cue appeared and only then increased until movement onset (**Fig 3C**).

**Figure 3:**
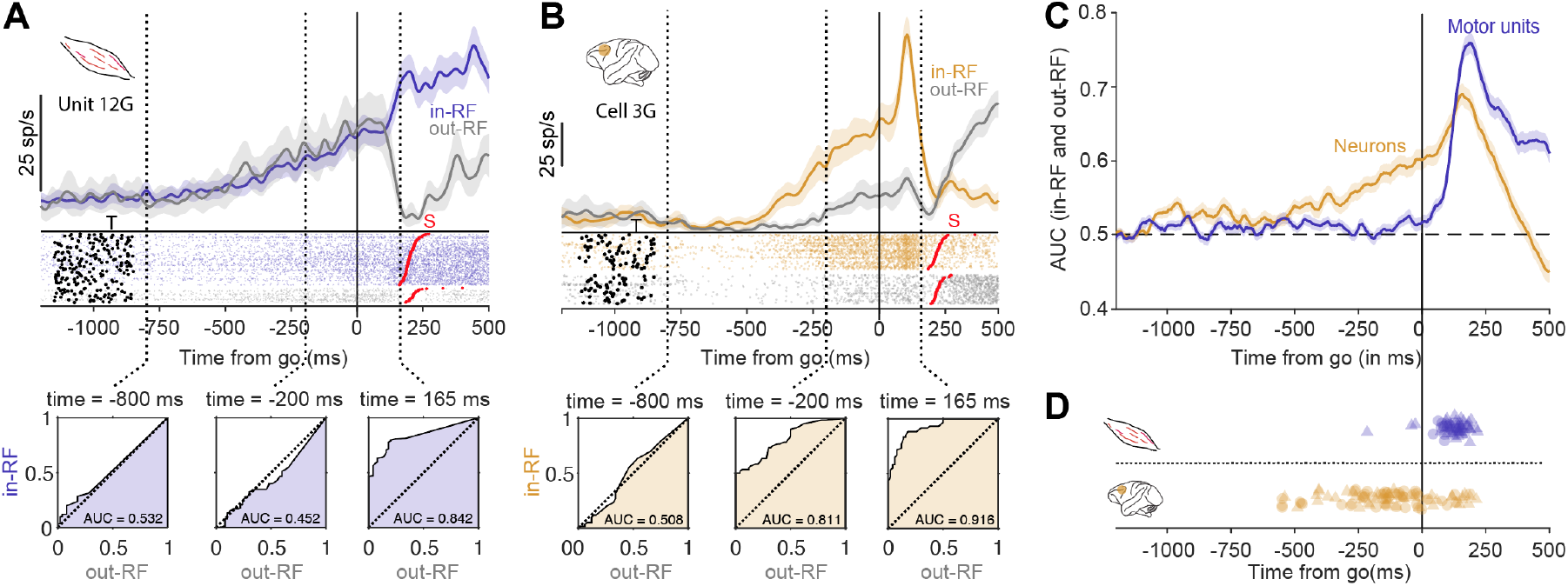
Spatial information about the saccade plan is present in the FEF neurons but not in the motor units. **A.** Top: a representative ramper motor unit with responses in-RF (purple) and out-RF (gray) conditions, aligned on the go cue. Raster, same format as **Fig 1F** . Vertical broken lines show the chosen time bins for ROC analysis. Bottom: ROC analysis for visual (left), delay (middle) and movement (right) time bin. **B.** Same as 3**A**, but for a representative neuron. **C.** AUC values from ROC performed between the in-RF and out-RF activities during the delay period for ramper motor units (purple) and neurons (gold). **D.** The direction discrimination time for ramper motor units (purple; top) and neurons (gold; bottom) for monkey J (circles) and monkey G (triangles).

We calculated the direction discrimination time, from the time of the go cue, for each unit that showed modulation during the delay period (**Fig 3D**). The average direction discrimination time for the entire population of spatially predictive neurons was −150.71 ± 25.47 ms for monkey J and −114.77 ± 30.80 ms for monkey G. The direction discrimination time occurred significantly before the go cue appeared for both monkeys (Monkey G: t(43) = −3.72; p <0.001; Monkey J: t(48) = −5.91; p <0.001). In contrast, for motor units, the average direction discrimination time was +104.66 ± 9.54 ms for monkey J and +128.78 ± 11.70 ms for monkey G. These onset times occurred significantly after the appearance of go cue (Monkey G: t(40) = 11.00; p<0.001; Monkey J: t(14) = 10.96; p<0.001).

These analyses indicate that although the direction of the upcoming saccade could be predicted by the activity of neurons during the delay time, motor units did not contain any information, despite being modulated during the delay period. That is, the spatial information encoded in the FEF in the delay period is not available in the peripheral musculature.

### Temporal information about the saccade plan during the delay period in the FEF neurons leaks into the neck muscle motor units

To test whether the ramper motor unit activity during the delay period could be used to explain the variability in the reaction time (RT), we calculated a Pearson’s correlation coefficient for each ramper motor unit and FEF neuron. Only the in-RF trials were used for further analyses (unless mentioned otherwise). We identified 10 (out of 70 motor) units (see **Fig 4A** for a representative motor unit) and 26 (out of 93) neurons (see **Fig 4B** for a representative neuron) that showed significant negative correlations between reaction time and change in activity during the delay period from the baseline (**Fig 4C**). Across the population, the mean correlation coefficient was significant for both motor units (r = −0.058 ± 0.017, t(69) = −3.504; p <0.001) and neurons (r = −0.052 ± 0.018; t(92) = −2.949; p=0.002; **Fig 4D**). We also performed a binomial test for both neurons and motor units and found the instances of negative correlations to be significant and above chance level across the population (motor units: F(47,70,.5) = 0.001 and neurons: F(62,93,.5) < 0.001).

**Figure 4:**
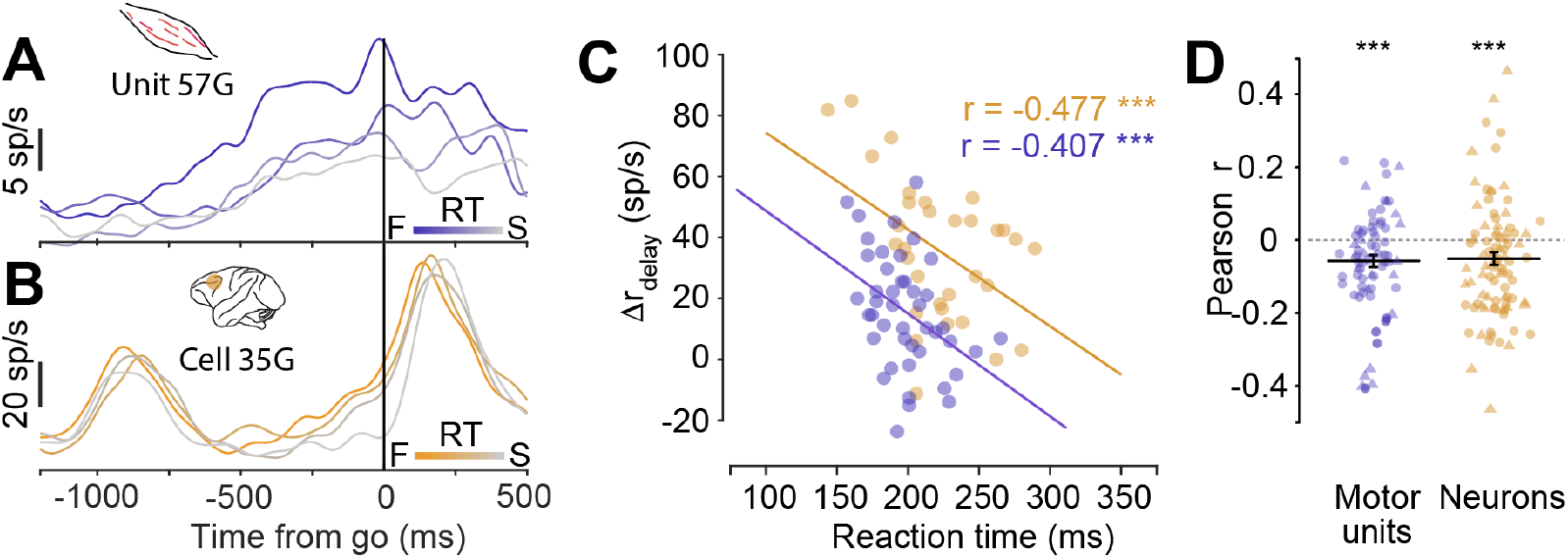
Temporal information about the saccade plan leaks from the center to periphery. **A.** A representative ramper motor unit’s activity aligned on the go cue for different reaction time (RT) values (see inset; F: fast, S: Slow). **B** . A representative neuron’s activity aligned on go cue for different reaction time values (see inset; F: fast, S: Slow). **C.** Scatter plots of EMG (purple) and neural activity (gold) during the delay period as a function of reaction time for a representative ramper motor unit and neuron respectively. **D.** Bee-swarm plot of Pearson’s correlations coefficient between activity in the delay period and reaction time for ramper motor units (purple) and neurons (gold). Circles and triangles represent data collected from monkey J and monkey G, respectively.

### Assessing the relationship between central and peripheral activity and behavioral reaction time

We tested whether activity observed during the delay time could be used to understand the roles of frontal eye fields and motor unit activity in determining reaction times. In previous studies (29) such ramping activity has been formalized in accumulator models which has been used only in the context of immediate movements to understand the neural basis of response preparation. Here, we tested if analogous parameters describing accumulation could be extended during the delay time to understand the activity of FEF neurons and motor units and their relation to reaction times. For each neuron and motor unit, we calculated different parameters of the accumulator model- (i) onset for the start of ramping activity during the delay period (ii) the growth rate of ramping and (iii) change in activity at the time of go cue, for slow and fast reaction time conditions (see methods).

The response of a representative ramper motor unit and a representative neuron for slow and fast reaction times are shown in **Fig 5A and 5C**. For both units, the activity around the go cue when compared to baseline, decreased with increase in reaction time (ramper motor unit: fast reaction time: 17.34 sp/s, slow reaction time: 9.90 sp/s; neuron: fast reaction time: 31.23 sp/s, slow reaction time: 16.89 sp/s). This observation was consistent across the population (ramper motor units: fast reaction time: 28.52 ± 1.82 sp/s, slow reaction time: 27.26 ± 1.86 sp/s; t-test: 43/70, t(69) = −2.50, p = 0.014; **Fig 5B**; and neurons: fast reaction time: 78.60 ± 5.60 sp/s, slow reaction time: 73.34 ± 5.59 sp/s; t-test: 71/93, t(92) = −4.63, p<0.001; **Fig 5D**). The change in activity at the time of go cue could be due to changes in the activity at onset or the rate of ramping during the delay time. More specifically, faster reaction times should be associated with larger rates and earlier onsets. For the example rasmper motor unit and neuron in **Fig 5A and 5C**, the change in activity at the time of go cue was due to systematic changes in the rate at which the activity rose during the delay period (ramper motor unit: fast reaction time = 15.50 sp/s, slow reaction time = 9.52 sp/s; neuron: fast reaction time: 83.94 sp/s, slow reaction time: 35.78 sp/s). Across the population, we observed significant changes in rate for both ramper motor units (fast reaction time: 24.97 ± 1.96 sp/s, slow reaction time: 21.24 ± 1.60 sp/s, t-test: t(69) = −2.51, p = 0.034; **Fig 5B**) as well as in neurons (fast reaction time: 70.38 ± 4.30 sp/s, slow reaction time: 60.46 ± 4.11 sp/s; t-test: t(92) = −3.47, p<0.001; **Fig 5D**, **Fig 5E**). However, we found no significant changes in onset values with increase in reaction time for ramper motor units for both fast reaction time (−637.98 ± 22.20 ms) and slow reaction time (−626.51 ± 22.28 ms) conditions (t-test: t(69) = 0.82, p = 0.41; **Fig 5B)** or neurons under fast reaction time (−534.13 ± 18.93 ms) and slow reaction time (−519.55 ± 18.85 ms) conditions (t-test: t(92) = 1.48, p = 0.14; **Fig 5D)**.

**Figure 5:**
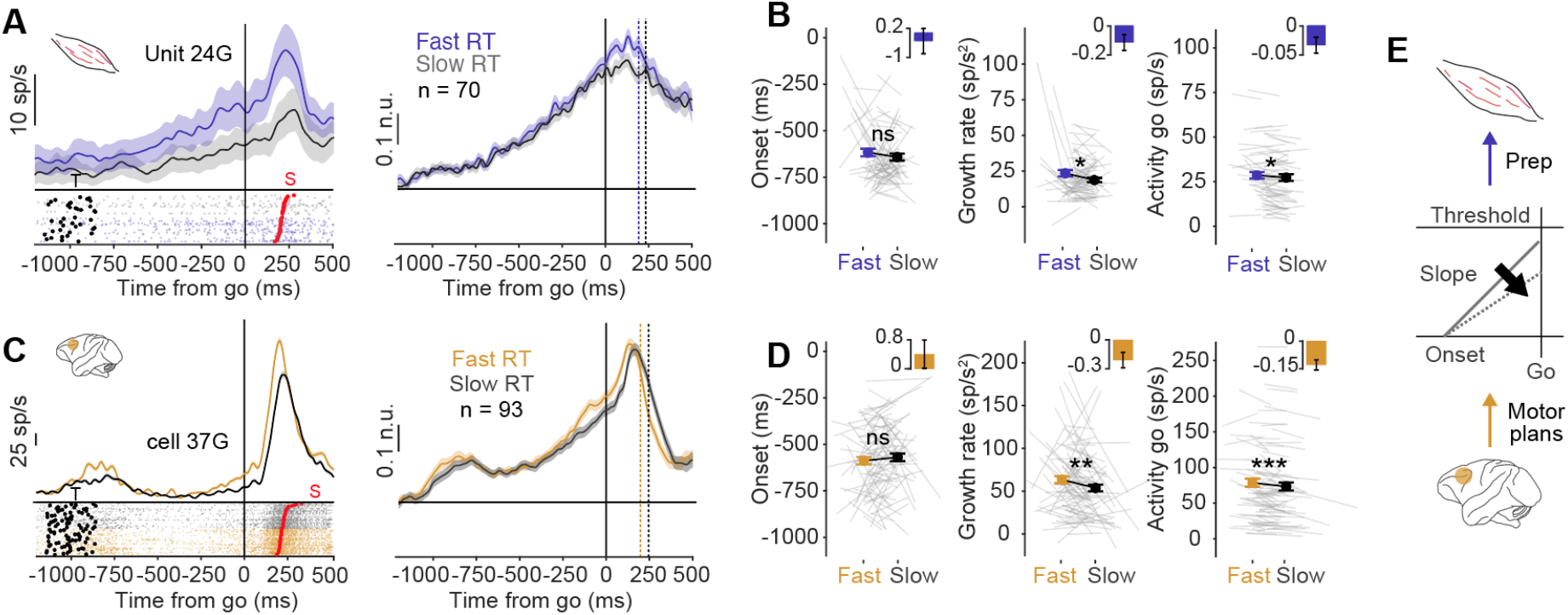
An accumulator model could explain the changes in activity at the center and periphery for slow and fast reaction times. **A.** Response of a representative motor motor unit (left; raster same format as **Fig 1F**) and the population response (right) aligned on the go cue for fast (purple) and slow (gray) reaction times. **B.** Accumulator model parameters for motor units: onset (left), growth rate (middle) and activity at go onset (right) for fast and slow reaction time conditions. The insets show the average and confidence interval for the slopes calculated for each line shown in grey, for the respective parameter. **C.** Same as **Fig 5A**, but for neurons. **D.** Same as **Fig 5B**, but for neurons. **E.** The accumulator model that best depicts the normalized activity for neurons and motor units.

Next, we also used delay time as a measure to understand the role of delay period activity in FEF and motor units in shaping reaction time. Since the target was displayed prior to the delay period, the monkeys could use the information to plan for an upcoming saccade. With a longer delay time, the monkeys had more time to prepare for their response as well as greater anticipation to expect the go cue. Thus, one of the benefits of having a delay time is that it leads to shorter reaction times, often called the delay time effect. Consistent with previous findings, reaction time was negatively correlated with the delay period (Linear Regression model: Monkey J: β=-0.008, p<0.05; Monkey G: β=-0.003, p<0.05, not shown in figure). We divided the response for each unit into short and long delay times. We also tested whether the same parameters could be used to explain the activity pattern of motor units and neurons using the same method described earlier.

For the representative ramper motor unit and neuron shown in **Fig 6A and 6C** respectively, the activity at the go cue was higher for longer delay periods (ramper motor unit: 32.68 sp/s; neuron: 68.24 sp/s) when compared to shorter delay periods (ramper motor unit: 23.91 sp/s; neuron: 56.51 sp/s). Both the units also showed a similar pattern of activity during the delay period. The rate at which the activity ramped up increased with increase in delay time (rate for motor unit at short delay time: 39.51 sp/s and long delay time = 40.81 sp/s; rate for neuron at short delay time: 106.57 sp/s and long delay time: 116.89 sp/s). Also, the ramping up started earlier for longer delay periods compared to the shorter delay period (onset for motor unit for short delay time: −630 ms and long delay time: −690 ms; onset for neuron for short delay time: −515 ms and long delay time: −580 ms relative to the go cue). Across the population, both the motor units and neurons showed significant differences in increase in the activity at the time of go cue from baseline with increase in delay time (activity for ramper motor units for short delay time: 25.76 ± 1.83 sp/s and long delay time: 29.77 ± 1.90 sp/s, t-test: t(69) =5.75, p<0.001 **Fig 6B;** activity for neurons for short delay time: 71.31 ± 5.51 sp/s and long delay time: 80.26 ± 5.81 sp/s; t-test: t(92) = 6.98, p<0.001 **Fig 6D**).We did not find and significant changes in growth rates with increase in delay time for motor units (rates for short delay time: 20.07 ± 1.54 sp/s and long delay time: 23.63 ± 1.97 sp/s; t-test: t(69)=1.97, p = 0.052; **Fig 6B**) but in the case of neurons, the population did show a tendency for rates to decrease with increase in delay time (rates for short delay time: 63.96 ± 4.21 sp/s and long delay time: 70.08 ± 4.84 sp/s; t-test: t(92) = 1.57, p = 0.41; **Fig 6D**). We found significant changes in onset for ramper motor units (onsets for short delay time: −630.24 ± 21.90 ms and long delay time: – 669.32 ± 23.42 ms, t-test: t(69) = −1.96, p = 0.027; **Fig 6B**) as well as neurons (onsets for short delay time: −497.77 ± 21.12 ms and long delay time: −568.44 ± 20.01 ms, t-test: t(92) = −3.88, p=0.001, **Fig 6D**).

**Figure 6:**
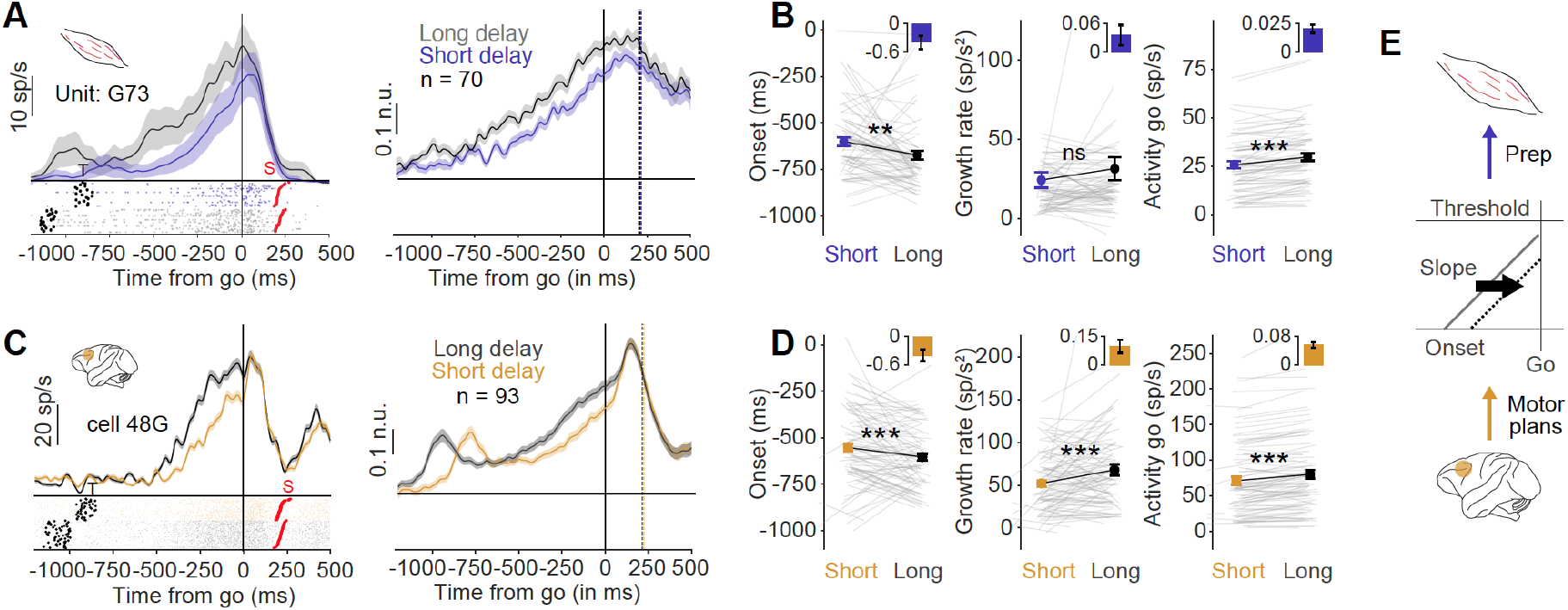
The same accumulator model could explain the changes in activity at the center and periphery for short and long delay times. **A.** Response of a representative motor unit (left; raster same format as **Fig 1F**) and the population response (right) aligned on the go cue for short (purple) and long (gray) delay times. **B.** Accumulator model parameters for motor units: onset (left), growth rate (middle) and activity at go onset (right) for short and long delay time conditions. The insets show the average and confidence interval for the slopes calculated for each line shown in grey, for the respective parameter. **C.** Same as **Fig 6A**, but for neurons. **D.** Same as **Fig 6B**, but for neurons. **E.** The accumulator model that best depicts the normalized activity for neurons and motor units.

In summary, by analyzing the data from two different perspectives-reaction time and delay time, similar conclusions could be drawn. Similar parameters could be used to explain modulations during the delay period in both FEF and neck muscles (**Fig 5E, Fig 6E**). Both, rate of ramping and onsets played an important role during the delay period to help initiate an upcoming saccade.

## Discussion

Our results show that motor units which were recruited early during the delay period had smaller amplitudes and represented the increased readiness to respond since their activity, like FEF neurons, was correlated with reaction time. Further, their activity profiles could be interpreted in a framework analogous to the accumulation models proposed to underlie reaction times: such that changes in the slopes and onsets could account for the slow and fast reaction times during the delay period providing a quantitative link between central and peripheral processing. We discuss these findings below.

### Relation to previous work

This work builds on previous work suggesting that unlike for eye movements, the neck muscles may not be as strongly inhibited by the omnipause neurons in the brainstem, due to the higher inertial of the head, thus allowing for gaze related commands in the FEF to be reflected rather early in their activity leading up to saccade and head movements (13). However, since gaze-related signals in neck muscles have not been studied in the context of delayed movements, the role of such activity in planning versus execution could not be ascertained until now. Compared to previous studies performed using gap paradigms and cued tasks (13, 15), we did not find many units that showed visually-evoked responses in the neck muscles. As noted before, such leakage of signals into the periphery may also be sensitive to context. In this regard, it is interesting to note that (21) did not observe any stimulus evoked responses in the human limb in a blocked delayed reach trials. However, subsequent work by Wood et al (30) showed that stimulus evoked responses can be seen in a delayed-reaching when it was intermixed with an immediate-reach condition. Thus, the context in which a cue is presented, and whether it may sometimes require an immediate response, may also be an additional factor for why stimulus-evoked responses were not observed in the current study. Additionally, the lack of stimulus-evoked responses may in part be consistent with the findings from (13), who showed that the stimulus-evoked response of the splenius capitis muscle was much lower, while deeper muscles like rectus capitis posterior and the obliquus capitis inferior muscles showed robust stimulus-locked responses. In this context, it should be mentioned that while we attempted to record primarily from the splenius capitis muscle, a large, superficial, ipsilateral head-turner, as it is easily accessible from the surface, the verification of the specificity of the penetration was not done. In addition to splenius capitis, the thickest layer and hence the other muscles most likely recorded from are the biventer cervicis/complexus muscles, which lie between the deepest layer rectus capitis muscle and the most superficial splenius capitis layer. The results suggest that our sample maybe consist of a mixture of splenius capitis and biventer cervicis/complexus muscles since the tuning curves of EMG activity to assess eye-EMG coupling showed a variety of responses ranging from preference for ipsilateral horizontal eye position/displacement, which is consistent with splenius capitis activity (13, 31–33), whereas units that showed preference for more upward movements may reflect biventer and complexus unit activity. However, since we did not find any consistent difference between the preferred direction of movement and the observed muscle activity during the delay period, we suggest that these properties are not muscle specific (**Fig S3**).

An additional difference between current work and previous work was that EMG signals were processed rather differently from conventional methods. As shown in **Fig 1E-G**, similar conclusions could be derived using either of the approaches. Since raster-based methods have been the standard approach to answer how information is represented in different brain areas, we decided to use the same approach to compare activity between the FEF and the neck motor unit activity in the periphery. This method allowed us to look at the recruitment process in more detail and revealed an expected heterogeneity amongst motor units allowing for us to distinguish and characterize rampers from non-rampers. Since non-rampers had action potentials that were typically smaller than the rampers, it is likely that they escaped detection in previous studies, with the EMG signal being dominated by the larger amplitude motor units that are more superficial and easily accessible (34). However, their smaller sizes may be a cause for concern given since the basal firing rates are higher than one would expect from an isolated motor unit. The average firing rates seem to be ~25sp/s, with some units exceeding 50 sp/s when aligned to saccade onset. When one considers that the animal is head-restrained in this experiment (and hence recruiting neck muscles as levels far below maximum voluntary contraction), the ramping activity we describe may likely be composed of multiple motor units. Nevertheless, despite this potential confound, the observation of distinct recruitment patterns suggests some degree of clustering of related motor units, which is corroborated by the well-defined movement fields observed for rampers and non-rampers alike (**Fig S3**). Finally, the presence of recruitment activity that predicts reaction time holds true independent of whether such activity is represented by single or multiple unit activity.

### Small motor unit recruitment and motor preparation

Our study shows the presence of heterogeneity in the motor units: While some units didn’t display any change in activity during the delay period (non-rampers), others did (rampers). What was however most interesting, and hitherto unreported, was that the rampers had smaller spiking amplitudes but larger spontaneous firing rates compared to the non-rampers. This finding is consistent with studies that indicate motor recruitment occurs according to Hennemann’s size principle in which smaller units, presumably coding for smaller forces are recruited first compared to larger motor units, presumably coding for larger forces (35). This hypothesis is consistent with the notion that smaller units had greater spontaneous firing rates and therefore are likely to have lesser inhibition, allowing for greater degree of transmission of information from the central gaze centers such as the FEF and SC. This finding is in line with the data by (36), who reported that the change in firing rate was larger with the slow conducting units and smaller with the fast conducting units that have smaller and larger recruitment thresholds, respectively.

Our findings parallel results obtained by Mellah et al (28) in monkeys performing a delayed reach movement. Similar to our study, they found the presence of two motor unit classes with the motor units associated with motor preparation (presumably rampers and slower fiber units coding for smaller forces) having low recruitment thresholds compared to the execution motor units that only responded during movement execution (presumably non-rampers and fast fiber units coding for larger forces). Interestingly, in accordance with our study, Mellah et al (28) also found that the preparatory motor units (rampers in our study) had smaller amplitude action potential compared to the execution motor units (non-rampers in our study). While the spike amplitude per se cannot be used as an absolute criterion to differentiate between the two classes, since it is known to depend on the distance from the electrode to the active fibre, the fibre diameter being the main factor determining the spike amplitude (37) and this might explain why the motor units recruited at low thresholds had lower amplitudes than those recruited at high thresholds (37–40). Thus, by analogy, the amplitudes observed in our experiments along with the discharge frequencies argue in favor of the idea that the motor units associated with motor preparation may be of the slow type, and that those associated with the execution may be of the fast type. Finally, it is also important to note that the presence of distinct classes we observe cannot be related to differences in the muscle types and other parameters that change across sessions as we observed these different classes of motor units and their associated recruitment patterns within a given session as well.

As suggested by Mellah et al, (28), the functional participation of each of these two types of motor unit in the preparation and execution of movements might be complementary. If the motor units associated with motor preparation are of the slow type, as seems to be the case, the tension developed by their activation might not suffice to initiate a movement. Slow motor units are known, however, to produce greater muscle stiffness than fast motor units subjected to similar tension levels (41, 42).Thus, it is possible that during the preparatory period, their activity may result in a certain level of muscle and head stiffness (since the muscle activity is not spatially specific) that amplifies the effect of the presumed fast motor units when the movement is triggered. The postural type of effects elicited by slow motor unit activity might thus tend to enhance the efficiency of the subsequent presumed fast motor unit discharge, allowing for rapid coordination between the eye and head movement despite differences in their inertia. Another function possibly served by the preparatory activation of the presumed slow motor units might be that of activating the proprioceptive and joint afferents which, by conveying inputs to the superior colliculus, contribute to building up the central nervous system activity responsible for the forthcoming movement; and more specifically, to controlling the excitability of neurons producing a phasic discharge which might activate the presumed fast motor units.

### Modeling a readiness signal in an accumulator framework in FEF neurons and motor unit activity

Our sample of FEF movement-related neurons showed the applicability of accumulation and the relation of specific parameters like onset, slope, activity level at the GO cue, and the ensuing reaction time, to the activity during the delay period. This observation extends previous work which has been limited to testing accumulation to threshold models relative to movement initiation after the GO cue. The presence of such activity prior to the GO cue indicates how prior expectation can modulate reaction times. Such integration is expected to occur earlier in the longer delay time conditions, thereby contributing to the shorter reaction times, which is what was observed in neurons and as well as ramping motor units. By recording from motor units in the neck muscle as a proxy of gaze-related preparation, we extended these observations to the peripheral musculature as well. However, unlike in FEF movement neurons that carry spatially specific signals during the delay period, only temporal information was observed with the rampers, which like their FEF counterparts was correlated with ensuing reaction times. The absence of spatial information in the rampers was striking and may reflect the task structure in which targets were 12 degrees in eccentricity which are unlikely to elicit head movements. Thus, the activity in motor units most likely reflects a strong saccade component and the absence of a head component may have produced a spatially non-specific signal. The non-specific nature of the rampers also rules out the possibility that this activity reflects neck muscle activity that was not executed due to head fixation. Instead, these data suggest that movement preparation or the readiness to respond is represented in the rampers which accounts for the correlations between motor unit activity during the delay time and reaction time. Such a temporal readiness signal is likely to reflect the outcome of the delay times being varied uniformly, by causing the expectation to increase with the delay time, and is likely to account for the delay period benefit in reaction time. This can lead to priming of the activity of muscles to actuate a movement.

A simple accumulator framework appears to fulfil the requirements of a unifying framework that could link a central process like movement preparation, to recruitment of motor units from the periphery, and behavioral reaction times. In this context, it is important to distinguish the accumulation that characterizes the subjects’ expectation or readiness to respond from the standard accumulator models of eye and hand reaction times, where accumulation to threshold describes the period between the GO signal and the movement (29, 43–47). In the accumulator framework, an increasing temporal expectation is actually modelled not as an increasing decision signal accumulation, but as an increased starting baseline (48). Therefore, the EMG ramp during the delay period might be evidence of how baseline increases (starting level after the GO cue) modulate reaction time. Since such temporal expectations are generated prior to the GO cue, and are expected to modulate or bias the reaction times, it is not surprising that we found low but significant correlations between measures such as onset, slope and activity both within neurons and rampers in relation to reaction time. However, what is interesting is that the relation between these measures remained the same for neurons and the rampers, indicating that the pattern of activity in motor units do reflect central processing that has bearing on behavioral reaction time.

## Acknowledgement

This study was supported by an Intensification of Research in High Priority Areas Grant from the Department of Science and Technology, Government of India; a Department of Biotechnology – Indian Institute of Science (DBT-IISc) partnership programme grant; and institutional support from the Ministry of Human Resource Development to A.M. S.P.R. and D.B. were supported by scholarship by Indian Institute of Science, Bangalore. The authors would also like to thank Prof. Brian Corneil for his inputs and suggestions, which has helped us to improve the quality of our work and manuscript.

## Conflict of Interest

The authors declare no competing financial interests

## Author Contributions

S.P.R., D.B. and A.M. designed the experiments, S.P.R. and D.B. performed the experiments, S.P.R.and N.S. analyzed the data, S.P.R.and N.S. made all the figures, all authors contributed to writing the manuscript.

## Methods

Some of the methods that were used in this study have been described in detail elsewhere (49–51). Here, we describe them briefly.

### Subjects

Two adult monkeys, J *(Macaca radiata,* male, age = 9 yrs; weight = 5.5 kg) and G *(Macaca mulatta,* female, age = 11 yrs; weight = 3.8 kg) were used for the experiments. All surgical procedures and monkey care were in compliance with the animal ethics guidelines of the Committee for the Purpose of Control and Supervision of Experiments on Animals (CPCSEA), Government of India, and the Institutional Animal Ethics Committee (IAEC) of the Indian Institute of Science and conformed to NIH (USA) guidelines as well.

### Memory-guided saccade task

Monkeys were trained to make a saccade to the remembered location of the target as shown in **Fig 1A**. After fixation on a central fixation point (FP: red, 0.6^0^ × 0.6^0^), for 300±15 ms, a target (T, grey, 1° × 1°) was flashed at the periphery of the screen for a duration of 100 ms. The target could appear at any one of eight locations lying on an imaginary circle with an eccentricity of 12^0^ from the fixation point. The monkeys had to maintain fixation until the fixation point disappeared. The disappearance of the fixation point was the ‘go cue’ to make a saccade to the remembered location of the target. The time interval or the delay period (1000 ±150 ms) between the go cue and the appearance of target, separated ‘where’ to look from ‘when’ to initiate a saccade. After a successful trial, the monkeys were given juice as a reward. A ±4^0^ and a ±6^0^ electronic window drawn from the center of the fixation point and the target, respectively, were used to assess the online performance of the two monkeys.

### Data collection

Experiments were carried out using a TEMPO/VIDEOSYNC system (Reflecting Computing, USA), simultaneously with Cerebus data acquisition system (Blackrock Microsystems, Salt Lake City, UT, USA). Eye positions were sampled with a monocular infrared pupil tracker (ISCAN, Woburn, MA USA), interfaced with TEMPO software real time. Visual stimuli were displayed using VIDEOSYNC software on a 42’’ LCD monitor with a refresh rate of 60 Hz and a resolution of 640 × 480 pixels. The monitor was placed 57 cm from the monkey sitting on a custom designed chair under head restrained conditions. While the monkeys were performing the task, we monitored their eye position and recorded simultaneous neural and intramuscular activity from the frontal eye fields and intermediate/deeper layers of neck muscles.

Raw neural signals from neurons were collected using single tungsten microelectrodes (FHC, Bowdoin, ME, USA; impedance: 2 to 4 MΩ) from the FEF on the right hemisphere through a permanently implanted recording chamber (Crist Instrument, Hagerstown, MD, USA). These raw neural signals were acquired at 30000 Hz and were subsampled to 1000 Hz and high pass filtered (300 to 3000 Hz) to obtain spikes.

Unilateral and bilateral intramuscular recordings were done using Polytetrafluoroethylene (PTFE)-coated stainless-steel needle electrodes of diameter 0.36 mm (TECA Elite series, Natus Neurology, USA) from dorsal neck muscles. Insertions were made with reference to externally available landmarks such as the occipital protuberance (horizontal) and the dorsal midline (vertical). All insertions were made beneath and within 2-4 cm of the horizontal and vertical reference landmarks, respectively. The collected EMG data was high pass filtered at 175 Hz, before further analysis. Neural and intramuscular and surface EMG recordings were analyzed in MATLAB with custom written software. A threshold criterion was used to extract units from the data set. The criterion (standard deviation = 2.25) was set based on the initial 500 ms of data from the beginning of each trial. The procedure was carried out using EMG LAB software (52). In short, the software collected the template of the waveforms (with window length of 2.33 ms for neurons and 8.33 ms for motor units) and the time stamps and for all the local maxima that were above the threshold were recorded as shown in **Fig 1**. Principal Component Analysis (PCA), template matching using a Euclidean distance method and correlations were used for sorting spikes. Initially, 3 to 4 templates were isolated for each trial using PCA. All the templates were then collected and based on these templates, further sorting was done. Typically, ~2-3 units were isolated by the experimenter, which were stable throughout the session (shown in **Fig 1**). Finally, based on the minimum least square distance value between each waveform and different templates, units were sorted into appropriate groups. The units which had similar amplitude size were clubbed together, irrespective of their phase. This was done because the activity of motor units is known to be sparse from previous studies and also to look at whether there is qualitative difference in response of motor units based on their size. Signal to noise ratio was calculated for each template using the method described in (53). The templates which had SNR values of less than 1.25 were rejected immediately. Some of the EMG templates which were very small in amplitude and had spike width less than 1.5 ms were rejected as noise. Based on visual inspection the spike width of motor units was greater than 3ms and had multiple phases. On the other hand, the neural data was much more well characterized and it was easier to detect noise owing to the spike width of single units, which were typically less than 1.5ms when compared to noise. Units were discarded if they could not be isolated clearly or if they did not show task-related response. After applying the exclusion criteria, a total of 210 neurons and 152 motor units were selected for further analysis.

## Data Analyses

The response of the units isolated from both neural and intramuscular recordings were analyzed in three epochs: visual epoch (50 to 250 ms after target onset); delay epoch (275 ms prior to 25 ms after the disappearance of fixation spot or go cue); and movement epoch (125 ms prior to 25 ms after saccade onset).

### Spike density function (SDF)

A spike density function was used to characterize the activity pattern of a neuron or motor unit. For each trial, the spike trains (of a unit) could be aligned with respect to the time when an event (example: onset of a saccade) occurred. The activity pattern of the unit could be estimated by averaging the convolved data over multiple trials. The discrete spike trains were convolved in both forward and backward directions using a Gaussian filter with a std. deviation of 10 ms. One advantage of processing signals using the above method is that all the features in the filtered data is exactly where they occurred in the unfiltered format.

### Response fields

The response field was defined for all those units which showed directional selectivity prior to a saccade. Out of the eight target locations, the four positions which showed higher activity during the movement epoch were grouped as inside response field (in-RF) and the diagonally opposite positions were grouped as outside response field (out-RF). A cosine tuning function was used to identify the preferred direction for each unit. The target locations which lied within ±90^0^ of the preferred direction were grouped as in-RF. The activity during the movement epoch for all correct trials was regressed with the direction of the target according to equation 1, using the Statistical toolbox in MATLAB.

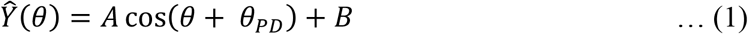

Here, Ŷ(θ) represents the activity of the unit when the target was presented at an angle θ°, B is the baseline firing activity of the unit and θ_PD_ is the preferred direction for the unit.

### Visuomotor Index

We classified FEF neurons into visual, visuomovement or movement units using a visuomotor index (VMI; equation 2) (54).

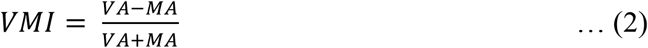

where VA is the visual activity in the visual epoch (90-180 ms after target onset) and MA is the movement activity in the saccade epoch (80 ms window preceding saccade onset). VMI ranges from +1 to −1. Visual neurons had positive VMIs, while movement neurons had negative VMIs and visuomovement neurons and intermediate VMIs. We only analysed visuomovement and movement neurons in this study. Their VMIs (mean ± s.e.m.) were – 0.203±0.032 and −0.301±0.054, respectively.

## Methods for analyzing EMG data

We used two methods to analyze EMG data. First, we used a standard method such as the root mean square of the EMG signal to analyze the recruitment of neck muscles. Mathematically, the root mean square measures the variability present in the signal and is thought to carry information about the recruitment of motor units in periphery. The raw EMG data was processed through 175 Hz low pass. For the filtered signal, a running window of length 35 ms with steps of 1ms was convolved with the square of the data. The square root of the convolved data for each trial was then used for further analysis. Additionally, we isolated spike waveforms from the EMG signal and extracted their time of occurrence and constructed a spike density function from this data. Both these approaches led to comparable results (**Fig 1;** Pearson’s r: 0.73; p<0.001).

To strengthen and support these findings, we also tested whether the smaller amplitude rampers might possibly reflect contamination due to noise crossing the threshold. However, there was no difference in signal to noise ratio (SNR) (Kelly et.al., 2007) between rampers and non-rampers (SNR_*Rampers*_: 2.406 ± 0.242, SNR_*Non--rampers*_: 2.5913 ± 0.1621; t(36) = −0.635, p = 0.53).

Furthermore, we used a standard ‘Additive time series decomposition model’ to split the activity of motor units into its underlying components, namely – trend, cycles and residuals (Equation 3, **Fig S1)**.

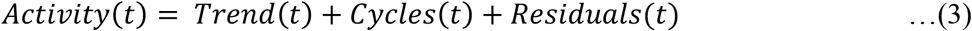

A moving average using a window of 30 bins or 150 ms was used to estimate the trend component present in the time series data. Followed by further decomposing the trend subtracted signal, into its cyclic components using the fast fourier transform method. Maximum of 2 to 3 peak cyclic components were identified as patterns and the remaining irregularity components were verified to be residuals or as white noise after using auto regressive and moving average (ARMA) approach. For this paper, we focus our analysis and discussions on underlying trend patterns for the time series data only.

### Direction discrimination time

For each unit, a standard receiver operator characteristic (ROC) analysis was performed on each time bin after the responses (in-RF and out-RF) were aligned on the go cue. The area under the ROC curve could range from 0.5 to 1. For each time bin, the ROC value of 0.5 would imply that the activity could not be reliably distinguished between in and out of the response field, whereas an ROC value of 1 would mean complete separation between the two distributions. A threshold criterion was set at 0.675 AUC, from – 450 ms to 350 ms aligned to the go cue. The first onset time at which the ROC value crossed the threshold for a continuous period of minimum of 25 ms prior to or from the go cue was marked as the direction discrimination time.

### Categorization of trials based on different parameters

Different parameter measures like reaction time and delay time were pooled across all the trials for all the neurons and motor units, recorded in each monkey separately. The pooled data for each parameter was then divided into two different groups. Based on this division, each trial was categorised into different conditions (i) slow and fast reaction times, (ii) short and long delay times for each unit recorded during saccadic eye movements.

### Components of the accumulator model

To test whether accumulator models could be extended during the delay period, various components of this framework were measured for each unit based on their spike density function, under in-RF conditions for (1) slow and fast reaction times and (2) short and long delay times. For each unit, activity at the go cue was measured by averaging the spiking activity 100 ms prior to and upto 50 ms from the go cue. Four different approaches were used to estimate the slope and onsets for capturing the trend component of each unit. First, a fixed interval from 900 ms prior to the go cue up to 25 ms was used to estimate the average slope using a simple linear regression approach. The time at which the global minima occurred over the same interval was considered as the onset value. Second, piecewise simple regression was carried out using a 250 ms moving window during the delay period to estimate the growth rate based on different metrics respectively-(a) maximum slope (b) best linear fit (r square). Under both the conditions, the value of the slope obtained at the global maxima index from the time series obtained after applying the moving window was considered as an estimate for the slope value. The estimated slope was then used to obtain the intercept on the x axis. Further, to be more conservative in our approach, the onset value was re-estimated to obtain the time at which the global minima occurred over the interval between and from the intercept value up to the go the cue. Third, a closing window approach with a rightward shift of 5 ms with each bin from 900 ms up to 25 ms prior to the go cue was used to estimate the growth rate using linear regression method. As described above in approach two, we used the slope value obtained at the global maxima index from the time series obtained in the previous step; to get an estimate for the slope value for both the metrics respectively. Similar to approach 2, the onset value was estimated to obtain the time at which the global minima occurred over the interval between and from the slope intercept on the x axis up to the go cue. Fourth, to be more conservative in our approach we combined piecewise regression and fixed interval method to estimate slope and onset values. In the first step, a piecewise approach was initially used to estimate the best fit slope and intercept on the x axis. Further, the onset value was re-estimated by regressing over the new fixed interval from step 1 up to 25 ms from the go cue. Finally, the global minima and slope obtained were considered as true estimates. Since the fourth method gave us the most consistent results, we used this method for all the analyses regarding accumulator models in **Figs 5 and 6.**

### Statistical tests

All statistical tests were done on combined data from both monkeys, unless specified otherwise. A Lilliefors test assessed the normality of the data under each condition. A n-way ANOVA was carried out on data when necessary, where n represents the number of multiple factors. A standard t-test or paired t-test was carried out whenever required. If normality failed, a sign rank or sum rank tests was done instead of paired and unpaired samples, respectively; and a Kruskal Wallis test was done instead of the ANOVA. A Binomial test was done to test whether the trend in the data was above chance level. In-built MATLAB functions and the statistical toolbox were used to carry out appropriate analysis.

---

**Figure S1.**
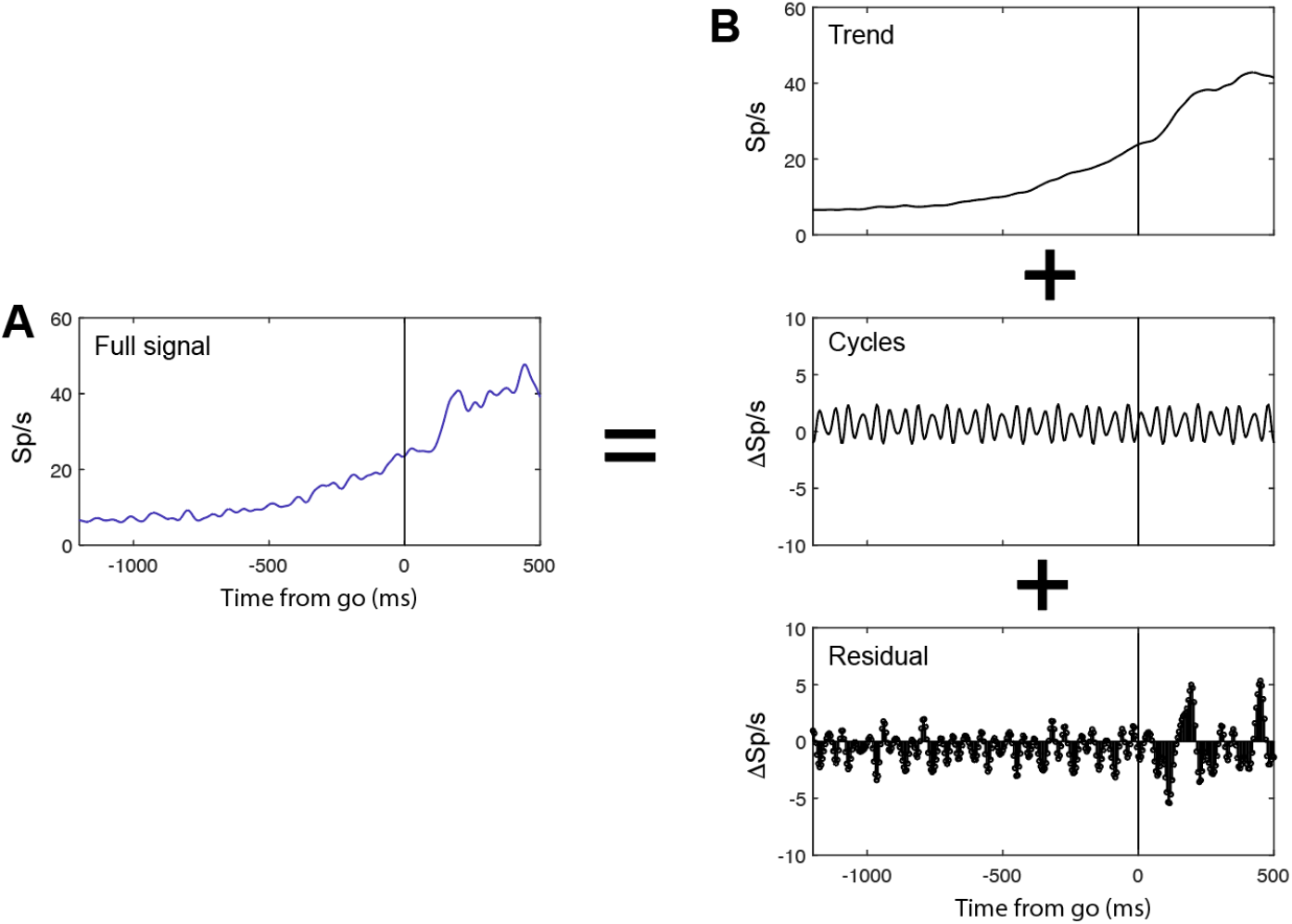
Trend, cycles and residual. **A.** Single unit activity (blue) aligned on go cue, for an upcoming saccade towards in-RF, for a representative motor unit recorded from neck muscle. **B.** Activity decomposed into individual components using an additive time series decomposition model – trend, cycles and residuals.

**Figure S2.**
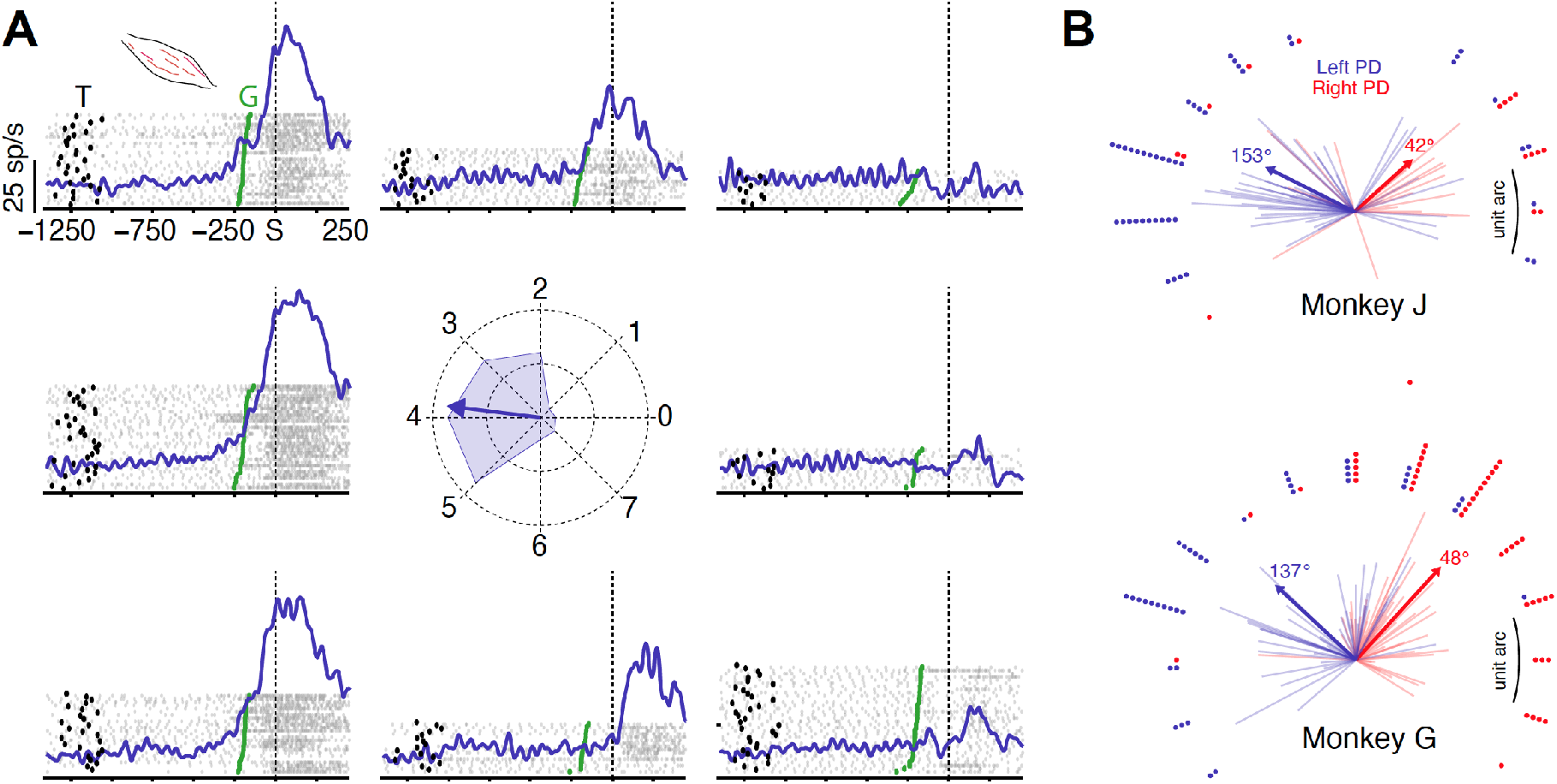
Motor units are spatially tuned. **A.** EMG responses (blue) aligned on saccade onset, for eight different target locations, of a representative motor unit recorded from the neck muscle. Each gray marker represents a spike. Thick black markers are time of target onsets. Each spike train represents the response on a single trial and the trials were sorted on the time of go cue (green markers). **B.** The plot at the center represents the preferred direction for the population (thick lines) and each motor unit (thin lines) that was recorded from left (blue) and right (red) neck muscles for monkey J (lighter shade) and monkey G (darker shade), respectively.

**Figure S3.**
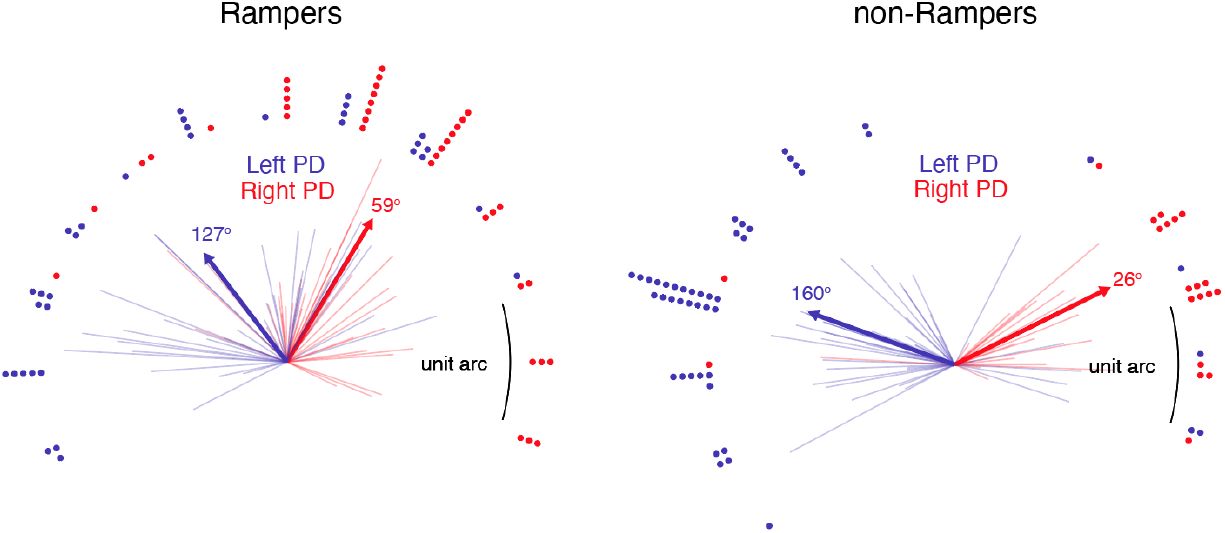
Preferred directions for ramper and non-ramper motor units. A plot of preferred direction for the population (thick lines) and each motor unit (thin lines) that was recorded from left (blue) and right (red) neck muscles for rompers (left panel) and non-rampers (right panel) from both the monkeys.

## Notes

### Competing Interest Statement

The authors have declared no competing interest.

